# Laboratory evaluation of the regeneration time, efficacy and wash-resistance of PermaNet® Dual (a deltamethrin-chlorfenapyr net) against susceptible and pyrethroid-resistant strains of *Anopheles gambiae*

**DOI:** 10.1101/2024.01.28.577674

**Authors:** Thomas Syme, Boris N’dombidjé, Damien Todjinou, Victoria Ariori, Corine Ngufor

## Abstract

Pyrethroid-chlorfenapyr nets have been recommended for malaria control by the World Health Organisation (WHO) after an alpha-cypermethrin-chlorfenapyr net showed improved impact in epidemiological trials. PermaNet® Dual is a new deltamethrin-chlorfenapyr net developed by Vestergaard Sàrl to expand options to control programmes. We performed a series of laboratory studies according to WHO guidelines to assess the regeneration time, efficacy and wash-resistance of PermaNet® Dual. Regeneration time was determined by subjecting net pieces to cone bioassays and tunnel tests before and 0, 1, 2, 3, 5 and 7 days after washing. The wash-resistance of PermaNet® Dual was evaluated compared to WHO-prequalified pyrethroid-only (PermaNet® 2.0) and pyrethroid-chlorfenapyr (Interceptor® G2) nets by testing net pieces washed 0, 1, 3, 5, 10, 15 and 20 times in cone bioassays and tunnel tests. Tests were performed with susceptible and pyrethroid-resistant strains of *Anopheles gambiae* to separately assess the pyrethroid and chlorfenapyr components. Net pieces were also analysed to determine insecticide content. In regeneration time studies, the biological activity of the deltamethrin and chlorfenapyr components of PermaNet® Dual regenerated within 1 day after washing and a 1-day washing interval was adopted for wash-resistance studies. PermaNet® Dual induced high mortality (98%) and blood-feeding inhibition (98%) of the susceptible strain after 20 washes fulfilling WHO efficacy criteria in tunnel tests (≥80% mortality, ≥90% blood-feeding inhibition). Similar results were obtained with PermaNet® 2.0 (99% mortality, 99% blood-feeding inhibition) and Interceptor® G2 (99% mortality, 98% blood-feeding inhibition) washed 20 times. In wash-resistance tunnel tests against the pyrethroid-resistant strain, PermaNet® Dual washed 20 times induced high mortality (91%) and blood-feeding inhibition (73%) which was similar to Interceptor® G2 (87% mortality, 79% blood-feeding inhibition) and superior to PermaNet® 2.0 (47% mortality, 68% blood-feeding inhibition). PermaNet® Dual fulfilled WHO efficacy criteria in laboratory bioassays and showed potential to improve control of pyrethroid-resistant malaria vectors.

## Background

The large-scale roll-out of insecticide-treated nets (ITNs) has been credited with most of the declines in malaria observed in sub-Saharan Africa over the past two decades (1). The public health value of ITNs is, in large part, attributable to the insecticide in the netting fibre, which kills or repels host-seeking vector mosquitoes providing protection against malaria for the user and the wider community. Pyrethroids have been the insecticide of choice on ITNs because they are highly effective, cheap, safe, and fast-acting (2). Overreliance on pyrethroids has however, selected for pyrethroid resistance in malaria vectors which is now pervasive throughout sub-Saharan Africa (3). Although the extent to which pyrethroid resistance is currently impacting the effectiveness of ITNs is unclear (4), malaria control progress has slowed on a global scale in recent years and in many high-burden countries downward trends in cases and deaths have reversed (5). Concern that pyrethroid resistance is contributing to stalling malaria control progress has stimulated substantial investment in development of new active ingredients (AIs) for use on ITNs.

Over the past decade, three types of ITN combining a pyrethroid with a second compound capable of improving control of pyrethroid-resistant malaria vectors have become available. The most promising next-generation ITN developed thus far are nets treated with the pyrrole insecticide chlorfenapyr (CFP). CFP is a new insecticide to vector control which induces mortality by uncoupling oxidative phosphorylation in the insect mitochondria (6). Because of its unique non-neurotoxic mode of action, CFP exhibits no cross-resistance with conventional neurotoxic insecticides and is hence a suitable AI to complement pyrethroids on ITNs (7). In laboratory and experimental hut studies, an alpha-cypermethrin-CFP net (Interceptor® G) improved mortality rates of pyrethroid-resistant malaria vectors to levels resembling that achieved with pyrethroid-only ITNs in areas of susceptibility (8–12). In subsequent epidemiological cluster-randomised controlled trials (cRCTs) in Benin (13) and Tanzania (14), Interceptor® G2 reduced child malaria incidence by 46% and 44% respectively over 2 years compared to pyrethroid-only ITNs and was found to be highly cost-effective. Based on the increasing body of evidence demonstrating the effectiveness and cost-effectiveness of pyrethroid-CFP ITNs, WHO has issued a strong recommendation for the distribution of pyrethroid-CFP ITNs over pyrethroid-only ITNs in areas of pyrethroid resistance (15). This is driving an increased demand for pyrethroid-CFP ITNs which are projected to comprise 80% of the African market by 2031 (16).

PermaNet® Dual is a new deltamethrin-CFP ITN developed by Vestergaard Sàrl that was recently added to the WHO list of prequalified vector control products (17) becoming available for large-scale deployment for vector control in endemic countries. To be prequalified by WHO, new ITNs are assessed for their safety, quality and entomological efficacy by the Prequalification Unit Vector Product Assessment Team (PQT/VCP). This includes laboratory studies demonstrating the efficacy and wash-resistance of the candidate ITN and the dynamics of the AI(s) in the netting fibre including the regeneration time (18, 19). In laboratory and experimental hut studies, the natural loss of insecticide under user conditions is simulated by washing. When the surface insecticide of a net is depleted after washing, it takes time for the reservoir insecticide in the netting fibre to migrate to the surface and restore full biological efficacy (20). The time taken for this to occur is the regeneration time which is used as the time interval between successive washes for wash-resistance studies. Adopting a wash-interval which allows full regeneration of insecticide at the net surface between washes is crucial to avoid generating biased results. If the regeneration time is too short for example, ITN durability may be overestimated because nets are washed before the insecticide has fully regenerated at the surface (21). Wash-resistance meanwhile is a critical predictor of the durability of an ITN as it indicates its ability to withstand washing and remain effective over several years of field use. It is assessed by subjecting nets to laboratory bioassays for up to 20 standardised washes which simulates insecticidal loss over 3 years under user conditions assuming nets are washed once every few months (19). Wash-resistance and regeneration time can be assessed entomologically in laboratory bioassays and chemically by measuring surface insecticide concentration on a net using reliable analytical methods.

We performed a laboratory study to evaluate the regeneration time, efficacy and wash-resistance of PermaNet® Dual following current WHO guidelines (19). The regeneration time of PermaNet® Dual was firstly assessed by subjecting net pieces to cone bioassays and tunnel tests unwashed and on successive days after washing. The wash-resistance of PermaNet® Dual compared to a WHO-prequalified pyrethroid-only ITN (PermaNet® 2.0) and pyrethroid-CFP ITN (Interceptor® G2) was subsequently evaluated by testing net pieces unwashed and after 1, 3, 5, 10, 15 and 20 washes in cone bioassays and tunnel tests. Regeneration time and wash-resistance studies were performed with susceptible and pyrethroid-resistant strains of *Anopheles gambiae* to separately assess the pyrethroid and CFP components of the ITNs. Net pieces were also analysed to determine within- and between-net variation and wash-resistance indexes of AI content. Following WHO PQT/VCP data requirements, the trial was performed in line with the Organisation for Economic Cooperation and Development (OECD) principles of good laboratory practice (GLP) at the CREC/LSHTM GLP-certified facility in Benin. This study predated the recently developed WHO guidelines for prequalification of ITNs (18) and was thus performed in accordance with the applicable guidelines for laboratory and field testing of ITNs previously published by WHO Pesticide Evaluation Scheme (WHOPES) (19). It was among the studies included in the WHO PQT/VCP prequalification assessment of PermaNet® Dual (22).

## Materials and methods

### Mosquito strain characterisation

Laboratory bioassays were performed with susceptible and pyrethroid-resistant strains of *An. gambiae* s.l. to separately assess the regeneration time and wash-resistance of the pyrethroid and CFP components of the ITNs. The rationale for this approach was two-fold. First, because pyrethroid-resistant mosquitoes were mostly expected to survive exposure to the pyrethroid, they could be used to quantify the effects of CFP. Second, because CFP is a slower-acting insecticide than pyrethroids taking up to 72 h to exert full toxicity, the mortality response of the susceptible strain after 24 h could be used to dissociate the effects of the pyrethroid from CFP and attribute activity to either compound.

The species composition and resistance profiles of the susceptible and pyrethroid-resistant strains used for the study are described below.

- *An. gambiae* sensu stricto (s.s.) Kisumu strain is an insecticide-susceptible reference strain originated from Kisumu, western Kenya.
- *An. gambiae* sensu lato (s.l.) Covè strain is an insecticide-resistant field strain which are F1 progeny of mosquitoes collected from CREC/LSHTM field station in Covè, southern Benin. Prior studies show that the strain exhibits a high frequency of resistance to pyrethroids and organochlorines but remains susceptible to other insecticide classes including CFP. Resistance is mediated by the knockdown resistance (*kdr*) L1014F mutation and overexpression of P450 enzymes, notably CYP6P3 (23).

### Susceptibility bioassays

The resistance status of mosquito strains can vary over time due to contamination events and/or changes in rearing conditions (24). We therefore performed susceptibility bioassays prior to the study to verify the resistance status of the *An. gambiae* s.s. Kisumu and *An. gambiae* s.l. Covè strains to the AIs in the ITNs. Approximately 100 mosquitoes of each strain were exposed to filter papers impregnated with the discriminating concentrations of alpha-cypermethrin (0.05%) and deltamethrin (0.05%) and bottles coated with the discriminating concentration of CFP (100 µg) for 60 mins in four cohorts of 20–25. Similar numbers of mosquitoes were concurrently exposed to silicone oil-impregnated papers and acetone-coated bottles as negative controls. After exposure, mosquitoes were transferred to untreated containers and provided access to 10% (w/v) glucose solution. Delayed mortality was then recorded after 24 h for alpha-cypermethrin and deltamethrin and every 24 h up to 72 h after exposure for CFP. Tests were performed at 27±2ᵒC and 75±10% relative humidity (RH).

### Net treatments and preparation of net pieces for bioassays and chemical analysis

We compared the efficacy and wash-resistance of the candidate ITN (PermaNet® Dual) to two other WHO-prequalified ITNs; a pyrethroid-only net (PermaNet® 2.0) and another pyrethroid-CFP net (Interceptor® G2). The technical specifications of the ITNs are described below.

- PermaNet® Dual (Vestergaard Sàrl) is a 100-denier, polyester ITN coated with a combination of deltamethrin and CFP at 2.1 g/kg and 5 g/kg respectively.
- PermaNet® 2.0 (Vestergaard Sàrl) is a 100-denier, polyester ITN coated with deltamethrin at 1.4 g/kg.
- Interceptor® G2 (BASF) is a 100-denier, polyester ITN coated with a combination of alpha-cypermethrin and CFP at 2.4 g/kg and 4.8 g/kg respectively.
- An untreated net developed to a similar technical specification to PermaNet® Dual was used as a negative control.

A total of four whole PermaNet® Dual, two PermaNet® 2.0, two Interceptor® G2 nets and one untreated control net were randomly selected to obtain net pieces for the laboratory bioassays and chemical analysis. The PermaNet Dual® nets were selected from three different production batches. Two sets of 14 net pieces measuring 25 x 25 cm were cut from each of the selected nets at WHO-recommended positions (19) (Figure 1). Five (5) net pieces from each of the PermaNet® Dual nets were set aside in a refrigerator at 4 °C for chemical analysis to determine within- and between-net variation in AI content. The remaining net pieces of all ITN types were then randomly assigned to study type (regeneration time, wash-resistance) and wash-point using sealed opaque envelopes. All net pieces were subsequently labelled, wrapped in aluminium foil and stored in an incubator at 30 °C before and between washes and testing. Net pieces were washed according to WHO guidelines (2) to deplete the surface availability of insecticide for both regeneration time and wash-resistance studies. To summarise, net pieces were placed in a 1 litre bottle containing standardised soap solution (Savon de Marseille dissolved in deionised water at 2 g/l) and washed for 10 mins in a shaker bath set at 155 movements per minute and 30 °C. Net pieces were then rinsed twice in clean deionised water under the same conditions. Following completion of the bioassays, all unwashed and washed net pieces used in regeneration time and wash-resistance studies were transferred to a refrigerator for storage at 4 °C before being sent for chemical analysis to determine the wash-resistance index of their respective AIs.

**Figure 1:**
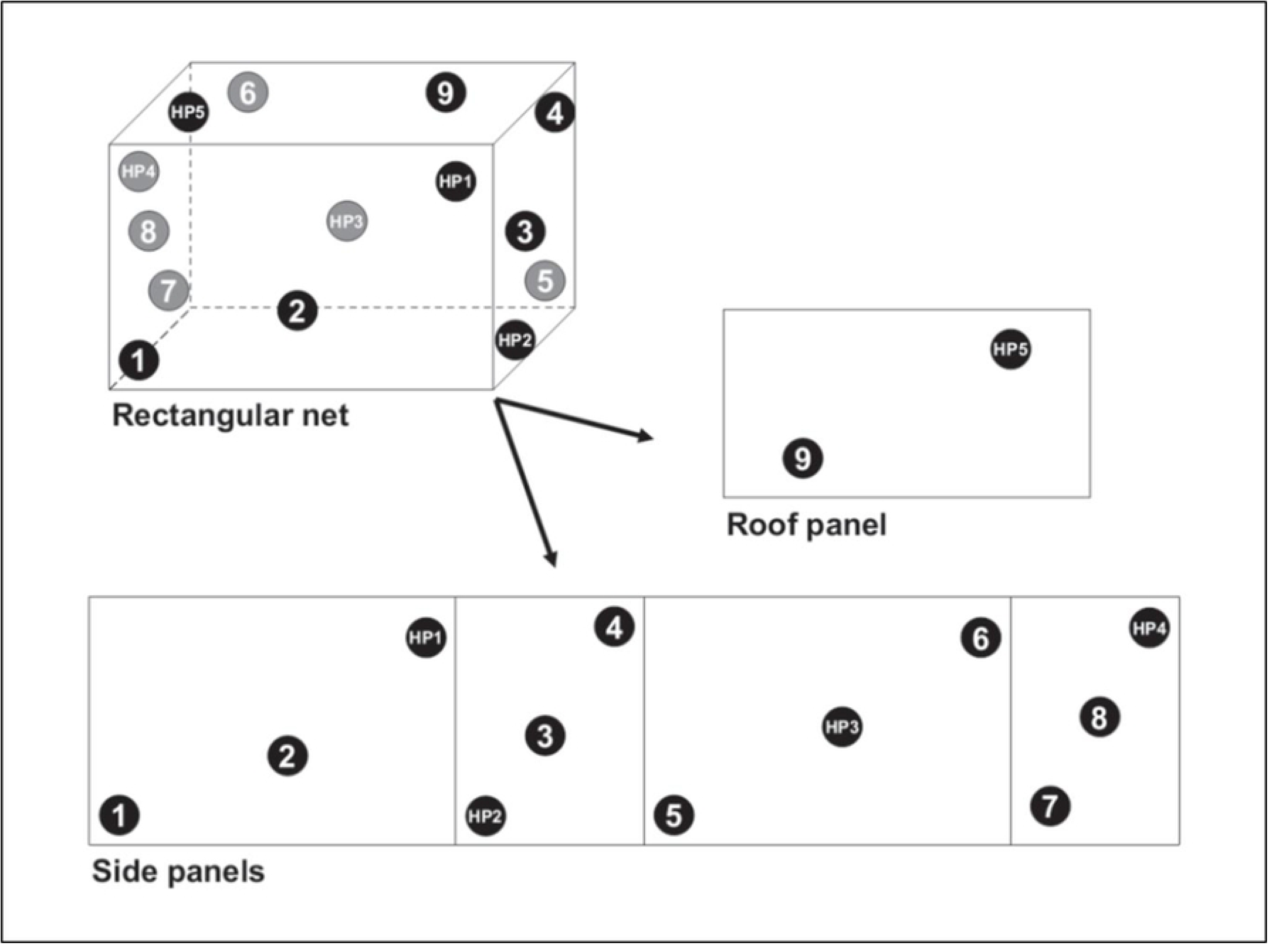
Sampling scheme for cutting 14 net pieces from whole nets for bioassays and chemical analysis. Source: World Health Organisation, 2013 (19).

### Laboratory bioassays

The regeneration time, efficacy and wash-resistance of PermaNet® Dual was assessed by performing WHO cone bioassays and tunnel tests on unwashed and washed net pieces against susceptible and pyrethroid-resistant mosquito strains under controlled laboratory conditions.

### Cone bioassays

The cone bioassay test design consists of plastic cones fixed to a frame containing a net piece. Mosquitoes aged 3–5 days were aspirated into the cones in cohorts of approximately 5 and exposed to the net piece for 3 mins. At the end of exposure, the mosquitoes were transferred to labelled holding cups and provided access to a cotton bud soaked in 10% (w/v) glucose solution. Knockdown was recorded 60 mins after exposure and delayed mortality every 24 h up to 72 h. Mosquitoes were concurrently exposed to untreated net pieces as a negative control. Tests were performed at 27±2ᵒC and 75±10% RH.

### Tunnel tests

Cone bioassays have been reported to underestimate the efficacy of nets treated with highly excitorepellent insecticides or AIs like CFP whose toxicity relies on the metabolic activity of the target insect (25). For this reason, ITNs failing to fulfil efficacy criteria in cone bioassays are subjected to tunnel tests in line with existing WHO guidelines (19). Tunnel tests are a type of experimental chamber which simulate the natural behavioural interactions that occur between free-flying mosquitoes and nets during host-seeking. The test design consists of a square glass tunnel divided at one third its length by a wooden frame fitted with a net piece which has been given 9 x 1 cm holes to facilitate entry in the baited chamber. In the short section of the tunnel, a guinea pig bait was held in an open-meshed cage while in the long section, approximately 100 mosquitoes aged 5–8 days were released at dusk and left overnight. In the morning, all mosquitoes were collected from the different sections of the tunnel and scored for immediate mortality and blood-feeding. Surviving mosquitoes were transferred to labelled holding cups and provided access to a cotton bud soaked in 10% (w/v) glucose solution after which delayed mortality was recorded every 24 h up to 72 h after exposure. Parallel exposures were performed with untreated net pieces as a negative control. Tests were performed at 27±2ᵒC and 75±10% RH.

### Preliminary assessment of test method and strain suitability

Despite high levels of pyrethroid resistance, laboratory-reared mosquito strains may not survive exposure to the pyrethroid component of nets in sufficient numbers to allow for assessment of the effects of the CFP component. Furthermore, previous studies have demonstrated the unsuitability of cone bioassays to assess the efficacy of CFP on ITNs (25). Hence, to validate our rationale and inform selection of the most appropriate methodology for regeneration time and wash-resistance studies, we performed a series of preliminary bioassays to compare the suitability of different test methods and mosquito strains for capturing the biological effects of the deltamethrin and CFP components of PermaNet® Dual. Mosquitoes of the susceptible *An. gambiae* s.s. Kisumu strain and pyrethroid-resistant *An. gambiae* s.l. Covè strain were exposed to new, unwashed net pieces of PermaNet® Dual, Interceptor® G2 and PermaNet® 2.0 in cone bioassays and tunnel tests. For cone bioassays, approximately 50 mosquitoes were exposed for 3 mins to each of four net pieces in ten cohorts of 4–6 while for tunnel tests, approximately 100 mosquitoes were exposed to each of two net pieces in one replicate tunnel. A total of approximately 200 mosquitoes per strain were thus exposed to each ITN type with both test methods. Parallel exposures were performed with untreated net pieces as a negative control.

### Regeneration time studies

Determining an accurate regeneration time is crucial for ITN evaluation as it defines the wash interval used for wash-resistance studies. To assess the regeneration time of PermaNet® Dual, we initially tested four net pieces unwashed in cone bioassays against the susceptible *An. gambiae* s.s. Kisumu strain and in tunnel tests against the pyrethroid-resistant *An. gambiae* s.l. Covè strain. The net pieces were then washed three consecutive times in the same day using a shaker bath as previously described and tested again in cone bioassays and tunnel tests against both strains 0, 1, 2, 3, 5 and 7 days after washing (Figure 2). On each day of testing, approximately 50 mosquitoes were exposed to each of four PermaNet® Dual pieces in ten cohorts of 4–6 in cone bioassays, while approximately 100 mosquitoes were exposed to each of two net PermaNet® Dual net pieces in one replicate tunnel test. A total of 200 mosquitoes were thus exposed to PermaNet® Dual pieces per time-point with both methods. The regeneration time was considered the time in days to reach a plateau in the biological efficacy after washing. Regeneration of the deltamethrin component of PermaNet® Dual was assessed based on knockdown after 60 mins and mortality after 24 h of the susceptible Kisumu strain in cone bioassays while regeneration of the CFP component was assessed based on mortality after 72 h of the pyrethroid-resistant Covè strain in tunnel tests.

**Figure 2:**
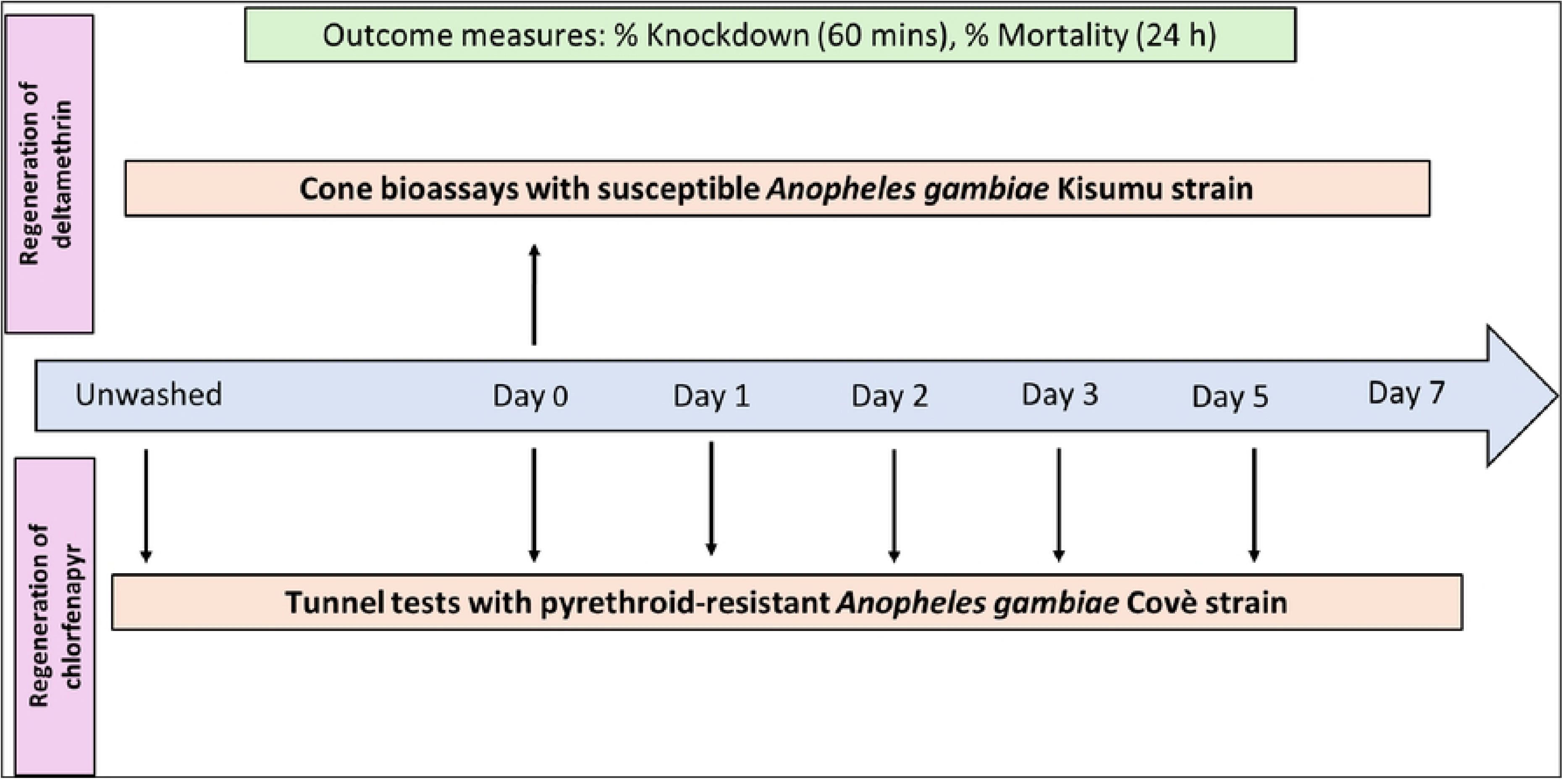
Testing scheme for regeneration time studies.

### Wash-resistance studies

In laboratory studies, a candidate ITN is expected to retain its biological efficacy after 20 standardised washes to fulfil WHO efficacy criteria (19). To assess the wash-resistance of PermaNet® Dual, we tested net pieces unwashed and washed up to 20 times in laboratory bioassays against susceptible and pyrethroid-resistant mosquito strains. Comparison was made to a WHO-prequalified pyrethroid-only ITN (PermaNet® 2.0) and pyrethroid-CFP ITN (Interceptor® G2) as positive controls. A total of four net pieces of each ITN type were washed in a shaker bath either 0, 1, 3, 5, 10, 15 or 20 times at intervals corresponding to the regeneration time as previously described. Net pieces were then tested in cone bioassays against the susceptible *An. gambiae* s.s. Kisumu strain and tunnel tests against the pyrethroid-resistant Covè strain to assess the wash-resistance of the deltamethrin and CFP components of PermaNet® Dual respectively. PermaNet® Dual initially failed to achieve WHO efficacy criteria in cone bioassays thus tunnel tests were also performed with the Kisumu strain to better assess the wash-resistance of its deltamethrin component. For cone bioassays, a total of approximately 50 mosquitoes were exposed to each of the four net pieces prepared per wash-point for 3 mins in ten cohorts of 5. Meanwhile for tunnel tests, approximately 100 mosquitoes were exposed to each of the two net pieces which were randomly selected from the cohort of four pieces washed 0, 10 and 20 times. A total of 200 mosquitoes were thus exposed per ITN and wash-point with both test methods. Efficacy criteria described in WHOPES guidelines were used as a benchmark to interpret the wash-resistance of PermaNet® Dual. Hence, PermaNet® Dual was considered to have retained its biological efficacy after 20 washes if it induced: ≥95% knockdown or ≥80% mortality in cone bioassays, or ≥80% mortality or ≥90% blood-feeding inhibition in tunnel tests.

### Chemical analysis of net pieces

Following completion of the bioassays, all unwashed and washed net pieces used in the regeneration time and wash-resistance studies were sent to the Centre Walloon de Recherches Agronomiques (CRA-W) in Belgium for chemical analysis to determine within- and between net-variation of the AIs in the ITNs and their respective wash-resistance indexes. Deltamethrin and CFP content in PermaNet® Dual and PermaNet® 2.0 pieces were extracted by sonication with heptane using dicyclohexyl phthalate as internal standard and determined by normal phase High Performance Liquid Chromatography with UV Diode Array Detection. Alpha-cypermethrin and CFP in Interceptor® G2 were extracted from net samples by sonication with heptane using dicyclohexyl phthalate as internal standard and determined by Gas Chromatography with Flame Ionisation Detection. Each method of analysis was performed using the internal standard calibration. The analytical methods used were based on validated and standardized methods published by the Collaborative International Pesticides Analytical Council (CIPAC). The chemical analysis results were used to calculate the proportional retention of active ingredients (AIs) after each wash for up to 20 standardised washes. The wash-resistance index of the ITNs was also calculated according to WHO guidelines (19) as follows:

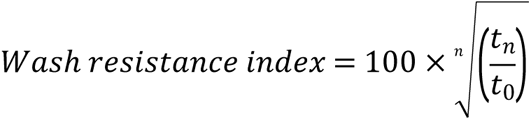

*Where: n = the number of washes, t_n_ = total AI content (g/kg) after n washes and, t_0_ = the total AI content (g/kg) with unwashed nets*.

### Data analysis

All laboratory bioassay data was recorded initially by hand on standardised data record forms before double entry into predesigned databases in Microsoft Excel. For regeneration time studies, proportional knockdown and mortality of mosquitoes were plotted against the number of days since washing. Knockdown and mortality of the Kisumu strain in cone bioassays was used to assess the regeneration time of the deltamethrin component of PermaNet® Dual while mortality of the Covè strain was used to assess the regeneration time of its CFP component. The time taken in days to reach a plateau in these outcomes was considered the regeneration time for each AI. For wash-resistance studies, knockdown and mortality for cone bioassays and mortality and blood-feeding inhibition for tunnel tests were plotted against the number of washes. As per the efficacy criteria outlined in WHOPES guidelines, PermaNet® Dual was considered to retain its biological efficacy after 20 standardised washes if it achieved ≥80% mortality and/or ≥95% knockdown in cone bioassays and ≥80% mortality and/or ≥90% blood-feeding inhibition in tunnel tests.

### Ethical considerations

We obtained ethical approval for the use of guinea pigs for blood-feeding of mosquito colonies and tunnel tests from the Animal Welfare Ethics Review Board of the London School of Hygiene & Tropical Medicine (LSHTM) (Ref: 2020-01). Guinea pig colonies were maintained at CREC/LSHTM animal house according to standard operating procedures (SOPs) developed in line with international regulations governing use of animals for scientific research purposes.

### Compliance with OECD principles of Good Laboratory Practice

To ensure compliance with the OECD principles of GLP, a series of activities were implemented during the initiation, execution, and reporting of the study. The study protocol was developed by a trained study director and approved by the sponsor before starting the study. Equipment used for the study (incubators and refrigerators for storage of ITN pieces, shaker baths for washing ITN pieces, and data loggers for monitoring ambient conditions) were calibrated before use. All ITN products used were verified to be within their expiry dates and were provided with certificates of analysis. The candidate PermaNet® Dual nets supplied by the manufacturer (Vestergaard Sàrl) were confirmed to come from three production batches. The environmental conditions under which the test products were stored were also verified daily using a calibrated data logger. Mosquitoes used for cone bioassays and tunnel tests were reared and transported in line with established SOPs that ensured the integrity of the strains tested. All computer systems (data loggers, databases, statistical software) used for data collection, entry and processing were validated before use. Physical records were kept of each procedure performed during the study. The quality assurance team of the CREC/LSHTM Facility performed inspections of the study protocol, critical phases of implementation, data quality and final report to assess compliance to GLP and no non-conformances were detected. The final report, along with all study-related documents, are securely stored in the physical and electronic archive of the Facility for up to 15 years. Study inspections performed in 2021 by the South African National Accreditation System (SANAS), the GLP certification body of the Facility, also detected no non-conformances.

## Results

### Susceptibility bioassay results

Mortality of the *An. gambiae* s.s. Kisumu strain was very high (≥98%) following exposure to the discriminating concentrations of alpha-cypermethrin, deltamethrin and CFP thus confirming its continued susceptibility to the AIs used in the ITNs (Table 1). In contrast, low proportions of the *An. gambiae* s.l. Covè strain were killed following exposure to the discriminating concentrations of alpha-cypermethrin (4%) and deltamethrin (10%) confirming that this strain continued to exhibit a high frequency of pyrethroid resistance. The discriminating concentration of CFP however induced 100% mortality of the Covè strain thus confirming continued susceptibility to the pyrrole. Mortality was negligible (≤4%) with the silicone oil and acetone controls with both strains.

**Table 1:**
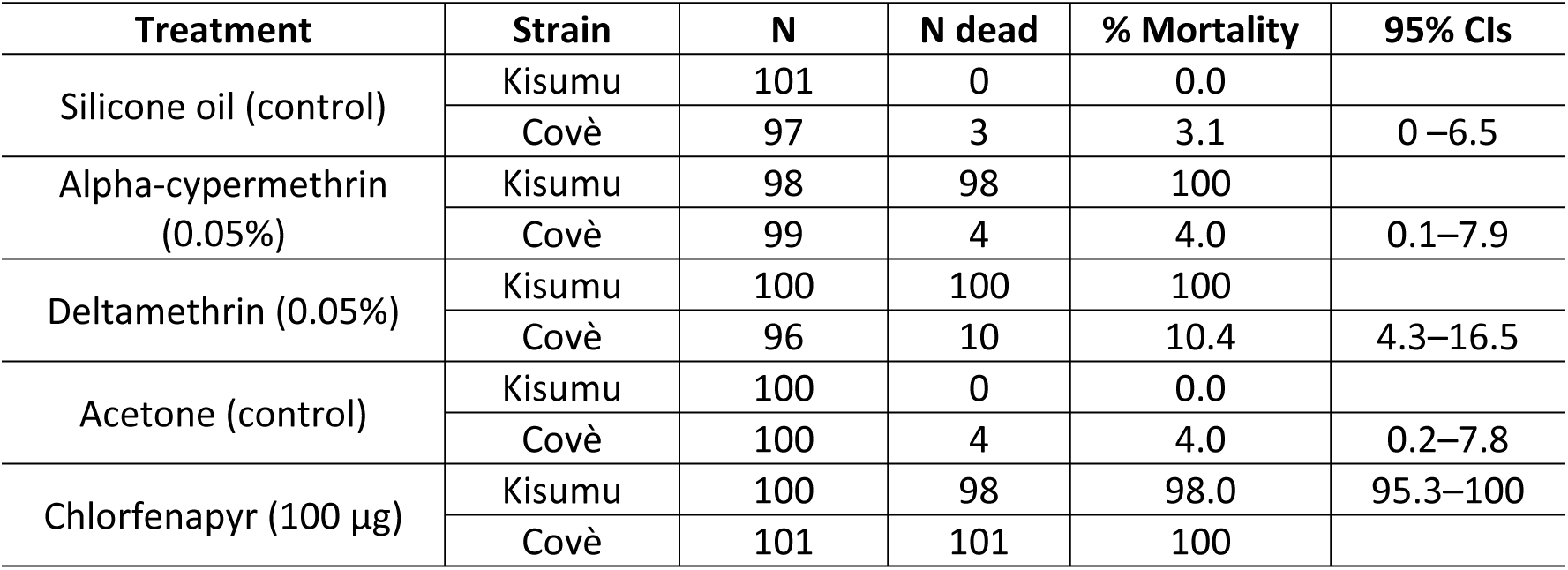
Susceptibility bioassay results with the Anopheles gambiae sensu stricto Kisumu strain and Anopheles gambiae sensu lato Covè strain. Approximately 100 mosquitoes of each strain were exposed to the discriminating concentrations of alpha-cypermethrin (0.05% filter paper), deltamethrin (0.05% filter paper) and chlorfenapyr (100 µg coated bottle) for 60 mins in four cohorts of 20–25. Mortality was recorded after 24 h for alpha-cypermethrin and deltamethrin while for chlorfenapyr, mortality was recorded after 72 h.

### Preliminary assessment of test method and strain suitability results

PermaNet® Dual induced moderate mortality (72%) of the susceptible Kisumu strain in cone bioassays while mortality rates were lower with Interceptor® G2 (33%) and PermaNet® 2.0 (44%) (Figure 3a). By contrast, all ITNs induced near maximum mortality (≥99%) of the Kisumu strain in tunnel tests (Figure 3b). Based on the moderate mortality response of the susceptible strain with PermaNet® Dual, cone bioassays were adopted as the primary method to assess deltamethrin on PermaNet® Dual with tunnel tests used only if efficacy criteria for knockdown (≥95%) and mortality (≥80%) in cone bioassays were not achieved (19). In cone bioassays performed against the pyrethroid-resistant Covè strain, mortality was low with both pyrethroid-CFP ITNs (7% with PermaNet® Dual and 27% with Interceptor® G2) and similar to the pyrethroid-only ITN PermaNet® 2.0 (10%). The low mortality rates observed with both pyrethroid-CFP ITNs in cone bioassays is attributable to the non-neurotoxic mode of action of CFP thus confirming the unsuitability of this method for assessing CFP on PermaNet® Dual. In tunnel tests however, both PermaNet® Dual and Interceptor® G2 induced near maximum mortality (∼99%) of the pyrethroid-resistant Covè strain. Based on this, tunnel tests were considered the more appropriate method for capturing the effect of CFP and were adopted for regeneration time and wash-resistance studies of CFP on PermaNet® Dual.

**Figure 3:**
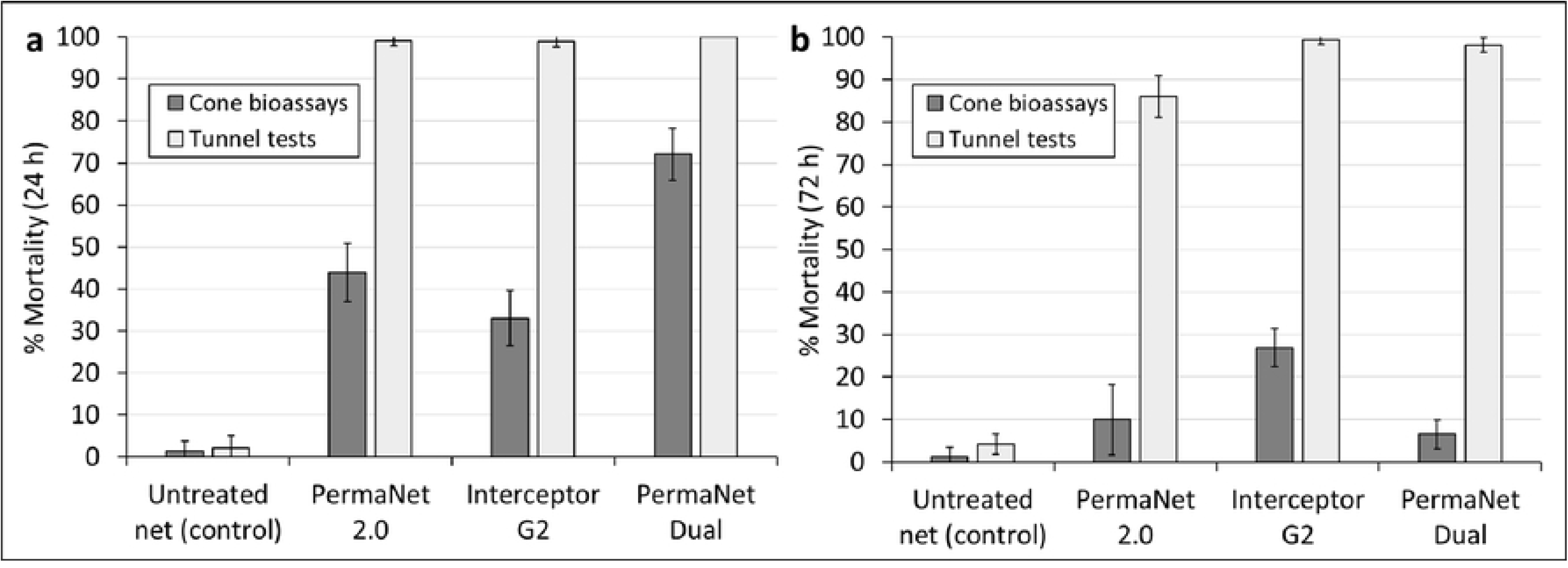
Mortality after 24 h of the susceptible *Anopheles gambiae sensu stricto* Kisumu strain (a) and mortality after 72 h of the pyrethroid-resistant *Anopheles gambiae sensu lato* Covè strain (b) in preliminary cone bioassays and tunnel tests to compare test methods and strain suitability. *A total of 200 mosquitoes per strain were exposed to each net type with both test methods. Error bars represent 95% confidence intervals*.

### Regeneration time results

#### Regeneration time cone bioassay results

Knockdown and mortality rates of the susceptible *An. gambiae* s.s. Kisumu strain in cone bioassays were initially used to assess the regeneration time of the deltamethrin component of PermaNet® Dual. Knockdown after 60 mins of the susceptible Kisumu strain was high (91%) following exposure to unwashed PermaNet® Dual pieces and was similar (88%) in cone bioassays performed with net pieces washed three times in the same day i.e. day 0 (Figure 4). In subsequent cone bioassays performed with washed net pieces at 1, 2, 3, 5 and 7 days after washing, knockdown remained consistently high (72–87%) showing no reduction after washing, with only slight decreases observed at days 2 (72%) and 3 (72%). A similar trend was observed with mortality. Unwashed net pieces induced 72% mortality after 24 h, and in cone bioassays performed on the same day with net pieces washed three consecutive times, mortality increased slightly to 79% (Figure 5). In subsequent cone bioassays performed with the washed net pieces at 1, 2, 3, 5 and 7 days after washing, mortality remained consistent (79–86%), showing no substantial differences. Knockdown and mortality remained within 10% after day 1 showing minimal changes in efficacy after this day and indicating establishment of a plateau from day 1. Knockdown and mortality with the untreated control net remained low (0–3%) throughout regeneration time cone bioassays. Based on these results, the knockdown and mortality effects deltamethrin in PermaNet® Dual against the susceptible Kisumu strain were judged to have regenerated in less than 1 day after washing. Detailed regeneration time cone bioassay results are provided in supplementary information (Table S1).

**Figure 4:**
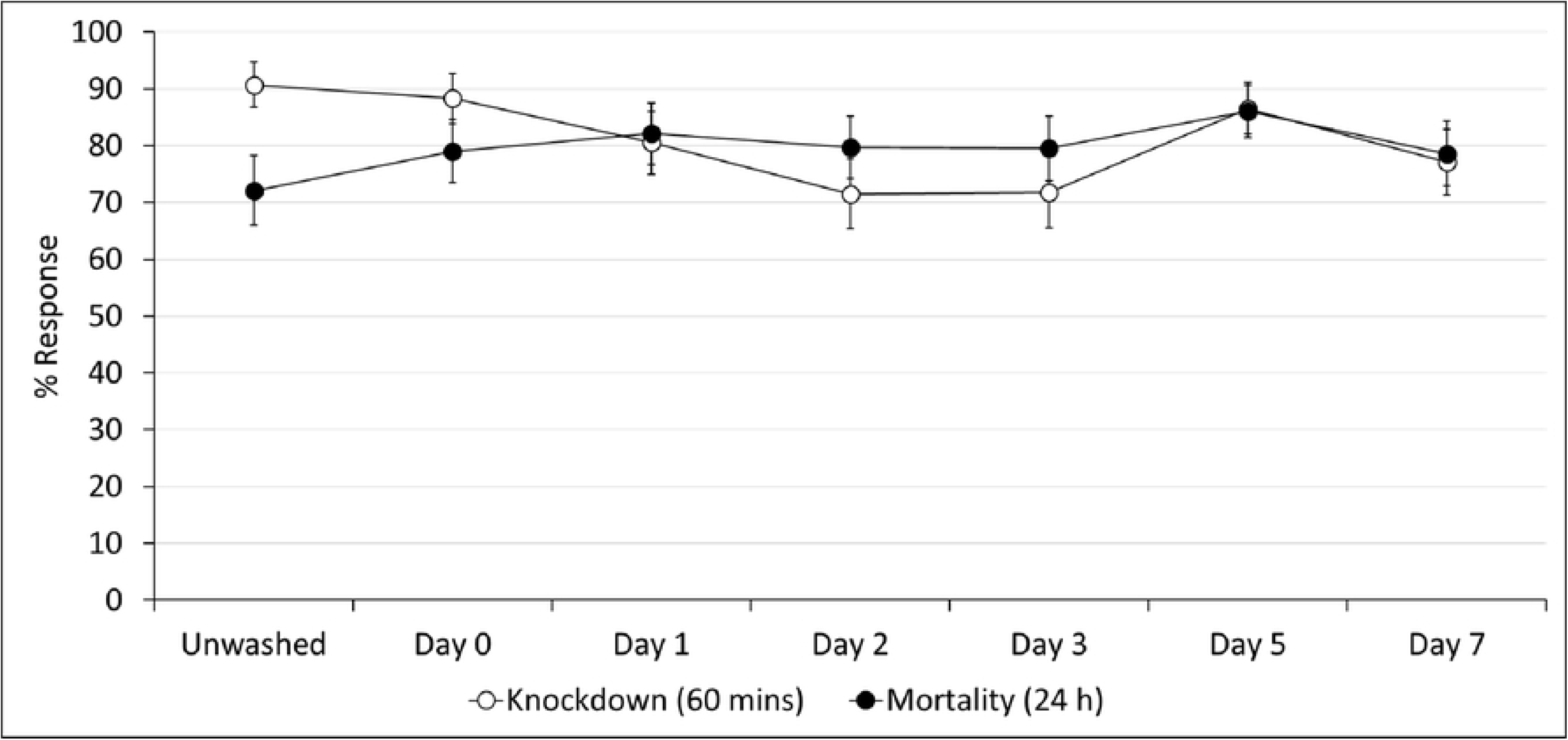
Knockdown after 60 mins and mortality after 24 h of the susceptible *Anopheles gambiae* sensu stricto Kisumu strain exposed to PermaNet® Dual net pieces in regeneration time cone bioassays. *At each timepoint, approximately 50 mosquitoes were exposed to each of the four unwashed and washed PermaNet® Dual net pieces for 3 mins in ten cohorts of 4–6. Error bars represent 95% confidence intervals*.

**Figure 5:**
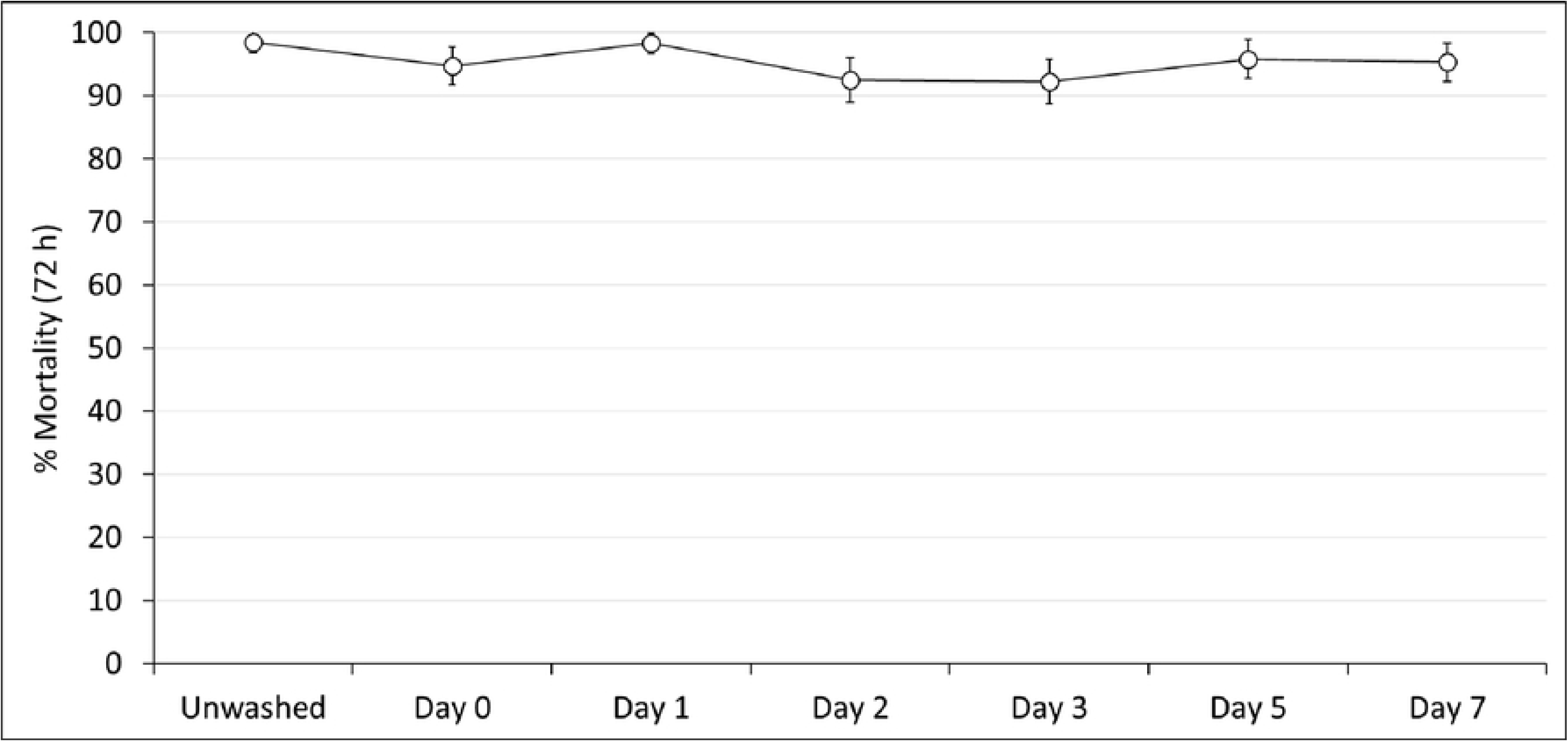
Mortality after 72 h of the pyrethroid-resistant *Anopheles gambiae sensu lato* Covè strain exposed to PermaNet® Dual net pieces regeneration time tunnel tests. *At each timepoint, approximately 100 mosquitoes were exposed to each of the two randomly selected unwashed and washed PermaNet® Dual net pieces overnight in one replicate tunnel test. Error bars represent 95% confidence intervals*.

### Regeneration time tunnel test results

Mortality rates of the pyrethroid-resistant Covè strain in tunnel tests were used to assess the regeneration time of the CFP component of PermaNet® Dual. Unwashed PermaNet® Dual net pieces induced very high mortality (98%) and in tunnel tests performed on the same day with net pieces washed three times, mortality was similar (95%) (Figure 5). In subsequent tunnel tests performed with the washed net pieces at 1, 2, 3, 5 and 7 days after washing, mortality remained consistently high (92– 98%). Mortality with the untreated control net was low (3–5%) throughout regeneration time tunnel tests. Based on these results, the mortality effect of chlorfenapyr in PermaNet® Dual against the pyrethroid-resistant Covè strain was judged to have regenerated within in less than 1 day after washing. A 1-day wash interval was thus adopted for wash-resistance testing of PermaNet® Dual. Detailed regeneration time tunnel test results are provided in supplementary information (Table S2).

### Wash-resistance results

#### Wash-resistance cone bioassay results

Knockdown and mortality rates of the susceptible Kisumu strain in cone bioassays were used to assess the wash-resistance of the deltamethrin component of PermaNet® Dual. As per the efficacy criteria outlined in WHO guidelines (19), the deltamethrin component of PermaNet® Dual was considered to have retained its biological efficacy after 20 washes if it induced ≥95% knockdown and/or ≥80% mortality in cone bioassays.

Unwashed PermaNet® Dual net pieces induced 78% knockdown after 60 mins of the susceptible Kisumu strain in cone bioassays and a similar knockdown effect was observed with washed net pieces ranging from 75–88% and failing to surpass 95% at all wash-points (Figure 6a). Mortality after 24 h with unwashed PermaNet® Dual net pieces was also low (39%) and although this increased at subsequent wash-points (46–76%), it remained <80% (Figure 6b). PermaNet® Dual thus failed to surpass WHO efficacy criteria for knockdown or mortality at any wash-point. With PermaNet® 2.0, knockdown was <95% at all wash-points tested meanwhile mortality was low with unwashed net pieces (44%) but increased to 66% after 1 wash and then to >80% at all subsequent wash-points except after 20 washes (72%). As with PermaNet® Dual, knockdown and mortality rates with Interceptor® G2 did not surpass WHO efficacy criteria at any wash-point. We observed no knockdown and very low mortality (2%) of susceptible Kisumu strain mosquitoes exposed to untreated control net pieces in wash-resistance cone bioassays.

**Figure 6:**
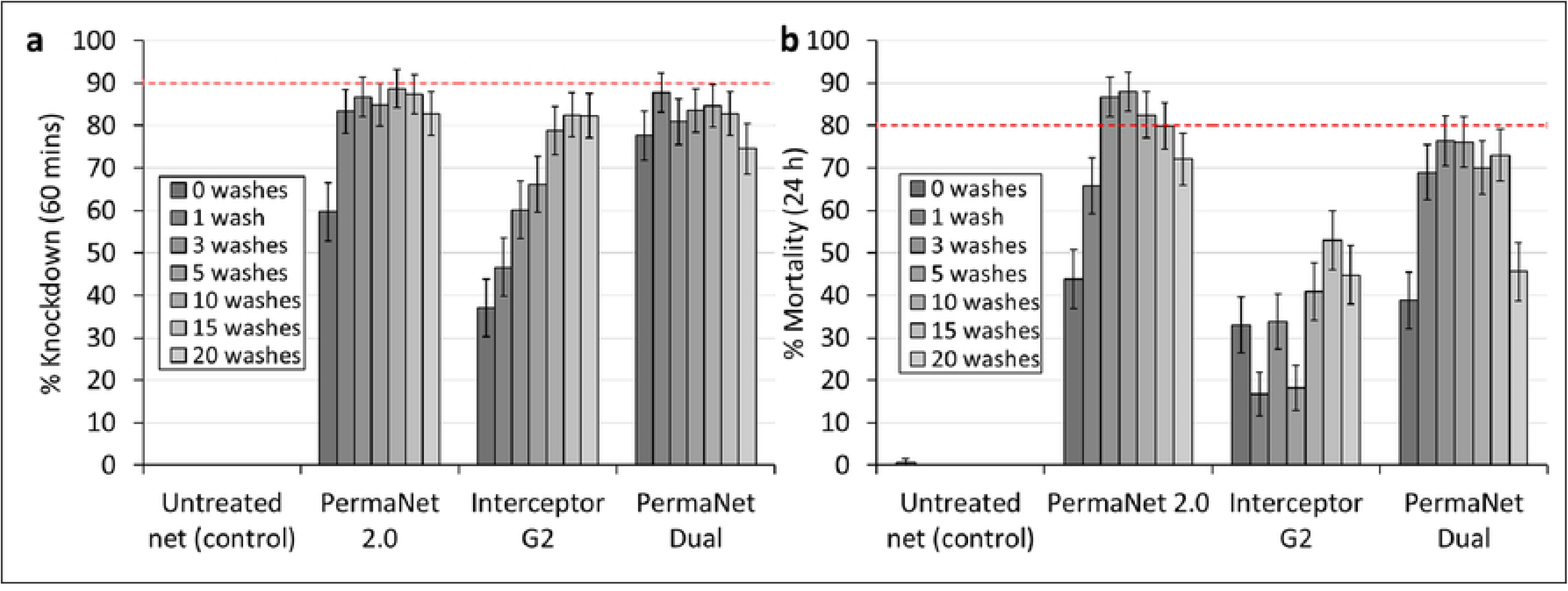
Knockdown after 60 mins (a) and mortality after 24 h (b) of the susceptible *Anopheles gambiae sensu stricto* Kisumu strain in wash-resistance cone bioassays. *At each wash-point, approximately 50 mosquitoes were exposed to each of the four net pieces of each net type for 3 mins in ten cohorts of 4–6. Red dashed lines represent efficacy thresholds for each outcome. Errors bars represent 95% confidence intervals*.

PermaNet® Dual thus failed to achieve WHO efficacy criteria for knockdown and mortality in cone bioassays against the susceptible Kisumu strain after 20 washes. WHO guidelines recommend that ITNs failing to achieve efficacy criteria in cone bioassays should be evaluated in tunnel tests. We therefore performed tunnel tests with the susceptible *An. gambiae* s.s. Kisumu strain to better assess the wash-resistance of the deltamethrin component of PermaNet® Dual. Detailed wash-resistance cone bioassay results are provided in supplementary information (Table S3).

### Wash-resistance tunnel test results

Owing to the failure of the candidate net to achieve WHO efficacy criteria against the susceptible Kisumu strain in cone bioassays, tunnel tests were performed with the same strain to better assess the wash-resistance of the deltamethrin component of PermaNet® Dual. Tunnel tests were also performed with the pyrethroid-resistant Covè strain to assess the wash-resistance of the CFP component of PermaNet® Dual and provide additional data on its efficacy against pyrethroid-resistant mosquitoes. Wash-resistance was assessed over separate time points for each AI i.e. 24 h for deltamethrin and 72 h for CFP which made it possible to dissociate and attribute activity to either compound. PermaNet® Dual was considered to have retained its biological efficacy after 20 washes if it induced ≥80% mortality and/or ≥90% blood-feeding inhibition in tunnel tests.

Unwashed PermaNet® Dual induced 100% mortality of the susceptible Kisumu strain and in tunnel tests with net pieces washed 10 and 20 times, mortality remained very high (≥98%) (Figure 7a). A similar trend was observed with PermaNet® 2.0 and Interceptor® G2, as almost maximum mortality (>99%) was observed with unwashed net pieces which remained very high (≥99%) after 10 and 20 washes. Unwashed PermaNet® Dual net pieces also induced high levels of blood-feeding inhibition (93%). In tunnel tests with net pieces washed 10 times, blood-feeding inhibition decreased slightly to 88% but, with net pieces washed 20 times this increased again to 98% (Figure 7b). Similarly, unwashed PermaNet® 2.0 and Interceptor® G2 net pieces induced 100% and 88% blood-feeding inhibition respectively, and these values remained very high (>98%) after 10 and 20 washes.

**Figure 7:**
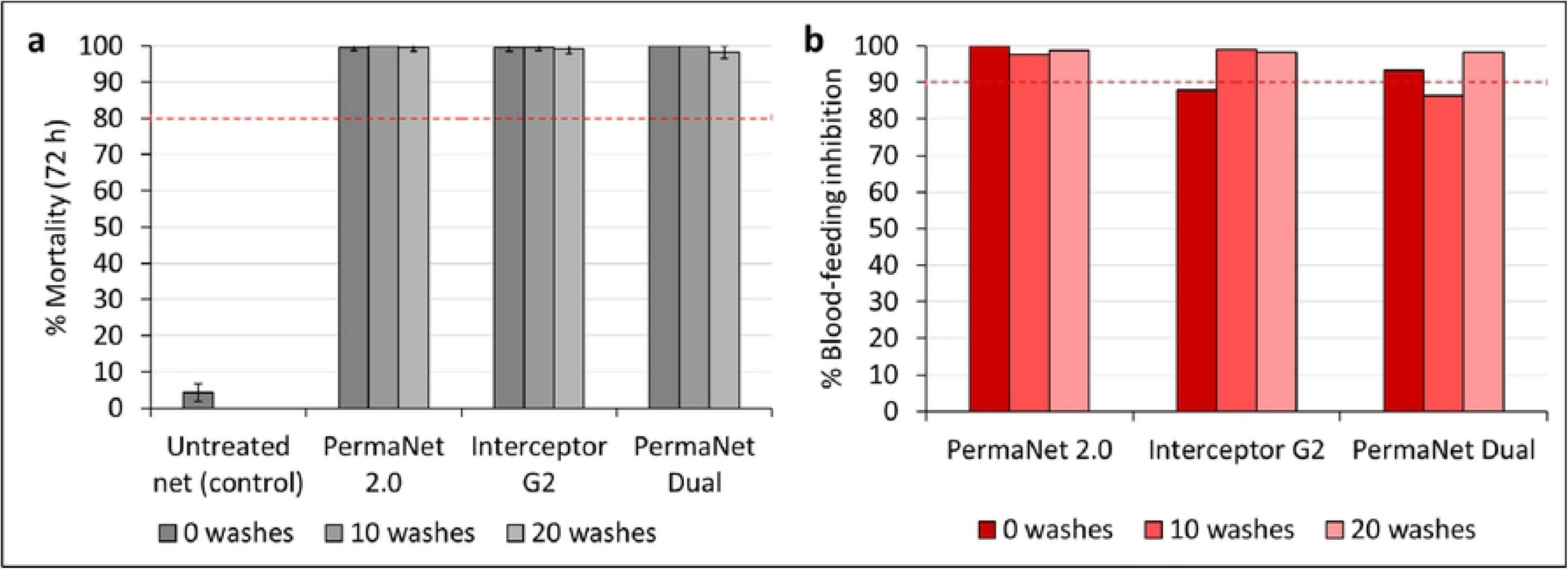
Mortality after 72 h (a) and blood-feeding inhibition (b) of the susceptible *Anopheles gambiae sensu stricto* Kisumu strain in wash-resistance tunnel tests. *At each wash-point, approximately 100 mosquitoes were exposed to each of the two net pieces of each net type overnight in one replicate tunnel test. Red-dashed lines represent efficacy thresholds for each outcome. Errors bars represent 95% confidence intervals*.

Unwashed PermaNet® Dual induced 98% mortality of the pyrethroid-resistant Covè strain and this remained very high with net pieces washed 10 times (97%) and 20 times (91%) (Figure 8a). A similar trend was observed with Interceptor® G2 which killed 99% of the Covè strain when unwashed and continued to induce high mortality rates after 10 (92%) and 20 washes (87%). In contrast, although we observed relatively high mortality of the pyrethroid-resistant Covè strain exposed to unwashed PermaNet® 2.0 pieces (84%), this fell considerably after 10 and 20 washes to 63% and 47% respectively. Blood-feeding inhibition was high with unwashed PermaNet® Dual pieces (93%) however, this decreased to 82% and 72% after 10 and 20 washes respectively (Figure 8b). A similar trend was observed with Interceptor® G2 and PermaNet® 2.0. Unwashed Interceptor® G2 and PermaNet® induced 93% and 92% blood-feeding inhibition respectively however, this fell below WHO cut-offs with net pieces washed 10 times (Interceptor® G2: 82%, PermaNet® 2.0: 84%) and 20 times (Interceptor® G2: 79%, PermaNet® 2.0: 68%). Both pyrethroid-CFP ITNs (PermaNet® Dual, Interceptor® G2) thus induced superior mortality and similar levels of blood-feeding inhibition against the pyrethroid-resistant Covè strain compared to the pyrethroid-only ITN (PermaNet® 2.0) demonstrating the potential of this net class to improve control of pyrethroid-resistant malaria vector populations.

**Figure 8:**
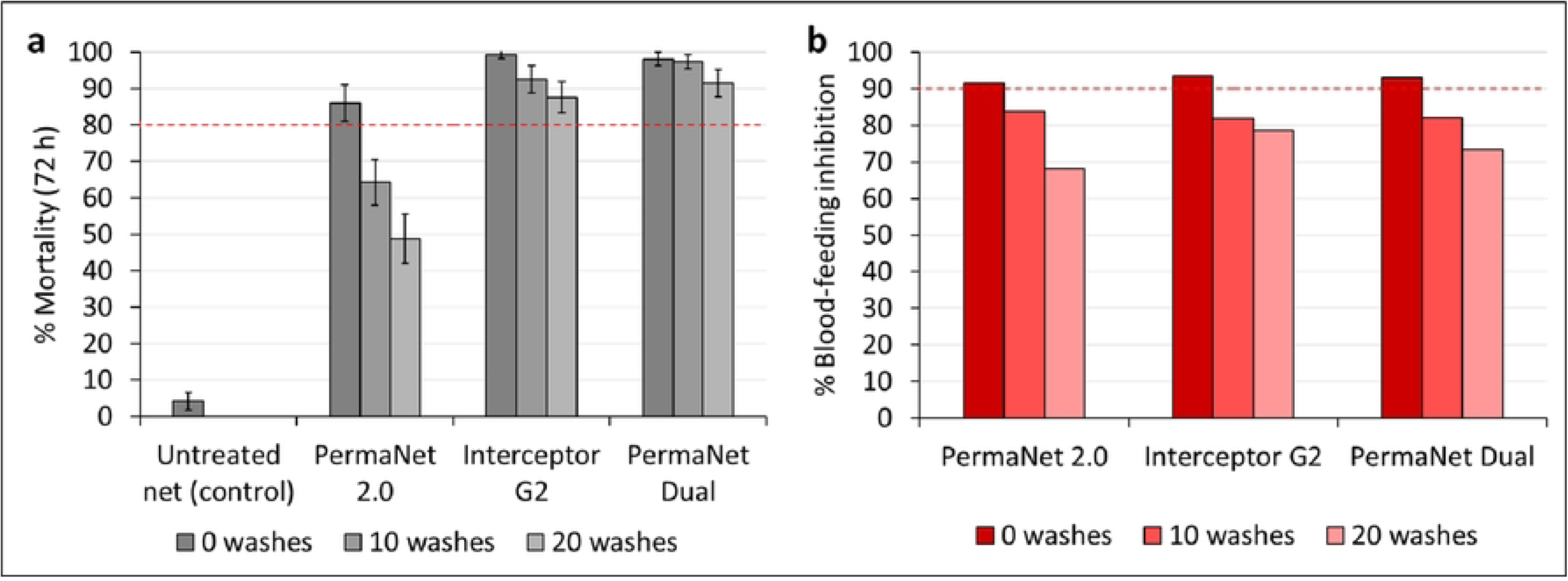
Mortality after 72 h (a) and blood-feeding inhibition (b) of the pyrethroid-resistant *Anopheles gambiae sensu lato* Covè strain in wash-resistance tunnel tests. *At each wash-point, approximately 100 mosquitoes were exposed to each of the two net pieces of each net type overnight in one replicate tunnel test. Red-dashed lines represent efficacy thresholds for each outcome. Error bars represent 95% confidence intervals*.

Mortality with mosquitoes exposed to untreated control net pieces was low (4%) for both strains while blood-feeding rates were high (76–79%). PermaNet® Dual thus achieved WHO efficacy criteria for mortality and blood-feeding inhibition against the susceptible Kisumu strain and for mortality against the pyrethroid-resistant Covè strain. Detailed wash-resistance tunnel test results are provided in supplementary information (Table S4).

### Chemical analysis of net pieces results

Analysis of the 20 PermaNet® Dual net pieces set aside for determination of within- and between-net variation revealed a mean AI content of 2.3 g/kg for deltamethrin and 5.1 g/kg for CFP showing that the nets complied with WHO tolerance thresholds (±25%) for both AIs (Table 2). The within-net variation expressed as the relative standard deviation (RSD) of the AI content in 5 net pieces obtained from the same net ranged from 1.9% to 15.0% for deltamethrin and 2.3% to 18.1% for CFP showing an acceptable homogeneity of both AIs on the net. The RSD of AI content in net pieces obtained from four different PermaNet® Dual nets was 3.3% for deltamethrin and 8.4% for CFP also showing an acceptable level of between-net variation.

**Table 2:** Within- and between-net variation of insecticide content in PermaNet® Dual net pieces. *\*Each value is the mean active ingredient content in 5 net pieces*.

| Net type | Active ingredient (AI) | Net no. | AI content (g/kg)* | Within-net variation % Relative standard deviation (RSD) | Mean AI content (g/kg) | Target AI content (g/kg) | % Difference | Between-net variation % RSD |
| --- | --- | --- | --- | --- | --- | --- | --- | --- |
| PermaNet Dual | Deltamethrin | 1 | 2.25 | 5.0 | 2.33 | 2.1 | +11.0 | 3.3 |
|  |  | 2 | 2.42 | 4.8 |  |  |  |  |
|  |  | 3 | 2.28 | 15.0 |  |  |  |  |
|  |  | 4 | 2.37 | 1.9 |  |  |  |  |
|  | Chlorfenapyr | 1 | 4.59 | 8.8 | 5.12 | 5.0 | +2.4 | 8.4 |
|  |  | 2 | 5.26 | 4.3 |  |  |  |  |
|  |  | 3 | 5.61 | 18.1 |  |  |  |  |
|  |  | 4 | 5.01 | 2.3 |  |  |  |  |

Analysis of net pieces washed 0, 1, 3, 5, 10, 15 and 20 times for wash-resistance studies showed that the deltamethrin component of PermaNet® Dual was more wash-resistant (24.2% retention after 20 washes) than CFP (5.4% retention after 20 washes) (Figures 9a and 9b). The wash-resistance index of deltamethrin was therefore higher than CFP after 20 washes (93.2% vs. 86.4%). Between the ITNs, there was a faster decline in deltamethrin content with PermaNet® 2.0 (11.8% retention after 20 washes) compared to PermaNet® Dual (24.2% retention after 20 washes). In contrast, there was a higher retention of AI content with Interceptor® G2 compared to PermaNet® Dual for both the pyrethroid (65.6% for alpha-cypermethrin vs. 24.2% for deltamethrin) and CFP components (29.7% vs. 5.4%) (Figures 9c and 9d). The wash-resistance index of deltamethrin after 20 washes was therefore higher with PermaNet® Dual than PermaNet® 2.0 (93.2% vs. 89.9%) while Interceptor® G2 had higher wash-resistance indexes for its pyrethroid and CFP components (97.9% with alpha-cypermethrin and

**Figure 9:**
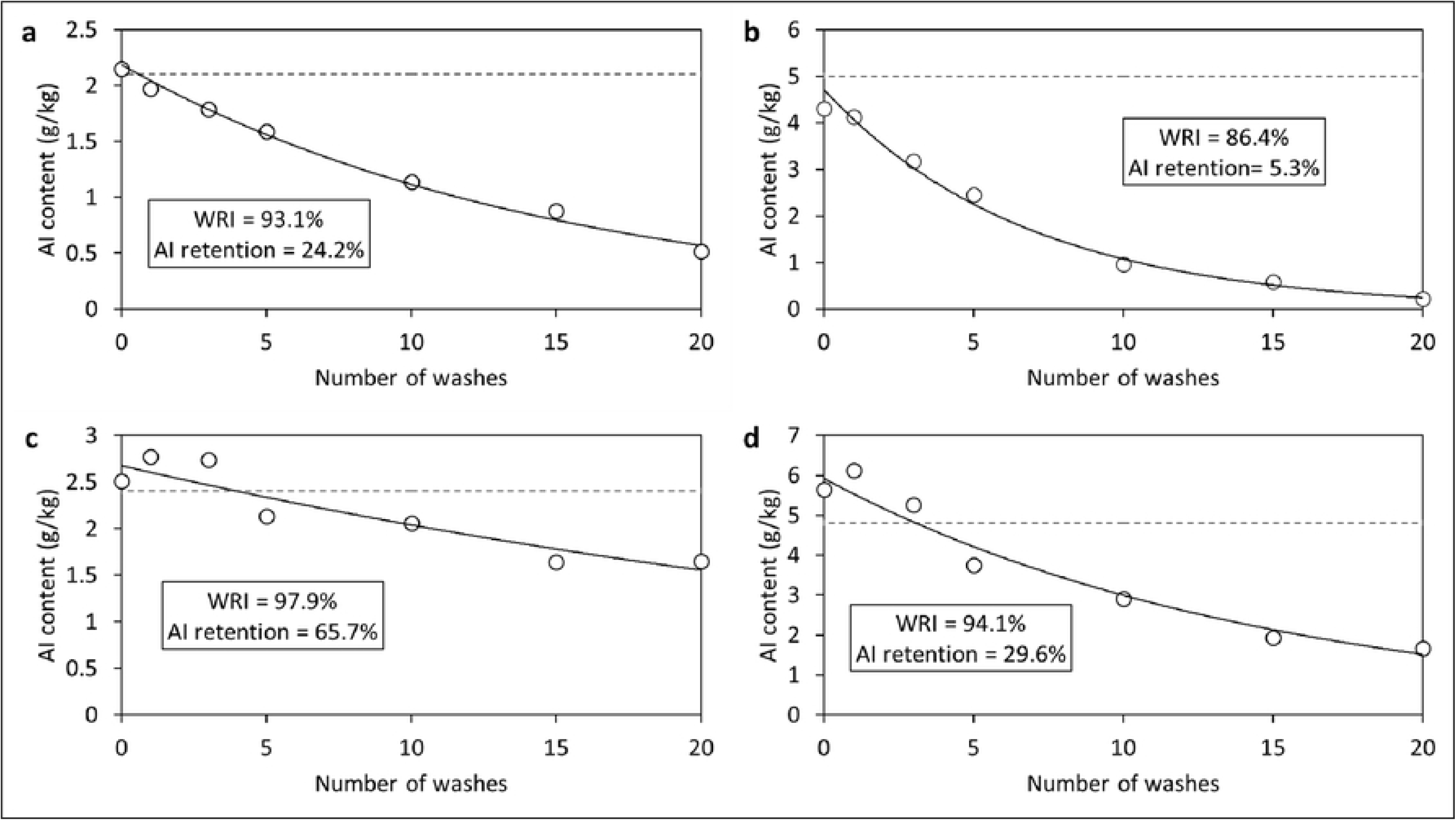
Wash retention of deltamethrin (a) and chlorfenapyr (b) in PermaNet® Dual and alpha-cypermethrin (c) and chlorfenapyr (d) in Interceptor® G2 net pieces used in wash-resistance studies. *Solid lines represent lines of best fit while dashed lines represent target active ingredient content. WRI=Wash-resistance index, AI=Active ingredient*.

94.1% with CFP) compared to PermaNet® Dual (93.2% with deltamethrin and 86.4% with CFP). Detailed chemical analysis results showing AI retention in nets after washing for regeneration time and wash-resistance studies are provided in supplementary information (Table S5).

## Discussion

To be added to the WHO list of prequalified vector control products and procured by major malaria control agencies, candidate ITN products must undergo a series of tests to demonstrate their safety, quality, and efficacy. These tests include laboratory studies to characterise the bioefficacy and physiochemical properties of the ITN. In this laboratory study, we assessed the efficacy, wash-resistance, and regeneration time of a new deltamethrin-CFP net (PermaNet® Dual) according to applicable WHO guidelines (19). PermaNet® Dual fulfilled WHO efficacy criteria against susceptible mosquitoes in laboratory bioassays and showed potential to improve control of pyrethroid-resistant malaria vectors.

The results showed that mosquito mortality with both pyrethroid-CFP ITNs was higher in tunnel tests than in cone bioassays, especially with pyrethroid-resistant mosquitoes as observed in the preliminary bioassays. Based on this, tunnel tests were used for studies assessing the regeneration time and wash-resistance of CFP on PermaNet® Dual. The unsuitability of cone bioassays for assessing the efficacy of CFP on ITNs is well-documented (25, 26). CFP is a pro-insecticide which owes its toxicity to the disruption of respiratory pathways in the insect mitochondria and because of this, it induces greater toxicity against insects with higher levels of metabolic activity (6). *Anopheles* mosquitoes exhibit peak flight activity during host-seeking which occurs at night due to the phase of their circadian rhythm

(27). Testing modalities such as the cone bioassay, which are conducted during the day and restrict mosquito flight activity, are therefore more likely to underestimate the efficacy of pyrethroid-CFP ITNs compared to overnight tunnel tests which simulate the normal behavioural interactions that occur between free-flying mosquitoes and nets during host-seeking. Our findings corroborate those of previous studies (25, 26) showing that tunnel tests with a highly pyrethroid-resistant mosquito strain represent a reliable method for assessing the activity of CFP on ITNs.

Laboratory and experimental hut studies assessing ITN efficacy simulate loss of insecticide by washing because this is considered among the primary drivers of insecticidal loss on ITNs under user conditions

(28). When surface insecticide is depleted after washing, the time taken for the reservoir insecticide to migrate to the surface and restore full biological efficacy – the regeneration time – is adopted as the wash-interval for these studies. Under real-life user conditions, nets are washed at intervals much greater than the regeneration time meaning they have enough time to regenerate between washes. For laboratory and experimental hut studies however, nets are washed at intervals corresponding to the regeneration time for practical reasons, as it represents the shortest possible time required for AIs to regenerate at the net surface after washing. In regeneration time studies, the biological activity of the deltamethrin and CFP components of PermaNet® Dual was consistently high showing no conspicuous dip in knockdown or mortality in the days after washing. This finding indicates that PermaNet® Dual may be a non-regenerating product. PermaNet® Dual is a coated net meaning the Ais are affixed onto the polymer matrix at the net surface in a fine layer of coating. This means that the AI does not need to migrate to the surface after washing and is thus immediately bioavailable to mosquitoes. The fast recovery of the biological activity of PermaNet® Dual after washing could therefore be attributable to its coating technology resulting in greater surface bioavailability of the AIs immediately after washing. By contrast, nets treated by the incorporation technology usually show slower recovery of biological activity after washing in regeneration time studies (17) because the AI is mixed into the polymer matrix of the net and would need to migrate to the surface of the net to become bioavailable to mosquitoes after washing. The rate of migration will also depend on multiple factors including the: size of the polymer matrix, types of polymers used for the coating, and ambient temperature (20).

Compared to Interceptor® G2, PermaNet® Dual induced higher knockdown and mortality of the susceptible Kisumu strain in cone bioassays and similar mortality of the pyrethroid-resistant Covè strain in tunnel tests. A review of the comparative efficacy of different types of pyrethroids showed that they induce broadly similar mortality responses of various laboratory-reared mosquito strains (29). Hence, the superior performance of PermaNet® Dual against the susceptible strain in cone bioassays was probably due to the higher loading dose and surface availability of pyrethroid in PermaNet® Dual rather than differences in the type of pyrethroid used on each net. The comparable performance of PermaNet® Dual and Interceptor® G2 against pyrethroid-resistant mosquitoes is consistent with a recent experimental hut trial in Benin which demonstrated the non-inferiority of PermaNet® Dual to Interceptor® G2 against wild, pyrethroid-resistant mosquitoes (30). Based on this and similar evidence of efficacy from trials in Côte d’Ivoire (31) and Kenya (22), PermaNet® Dual was added to the WHO list of prequalified vector control products (17) and included in the recent policy recommendation for pyrethroid-CFP ITNs (22). The findings of the present study reaffirm that PermaNet® Dual performs similarly to the first-in-class pyrethroid-CFP ITN Interceptor® G2 and represents an additional option of this highly effective net class for control of malaria transmitted by pyrethroid-resistant mosquitoes.

In laboratory studies, the ability of an ITN to remain effective over several years is assessed by subjecting nets to bioassays and chemical analysis for up to 20 standardised washes – a surrogate for insecticidal loss over 3 years of field use assuming nets are washed every few months (19). Both pyrethroid-CFP ITNs met WHO efficacy thresholds in tunnel tests after 20 washes corroborating previous studies (22, 30, 31) and thus showing their potential to remain efficacious over several years of household use. This suggests that the lower total CFP content retention observed with PermaNet® Dual compared to Interceptor® G2 had little or no impact on the durability of its insecticidal activity. An assessment of the amount of bioavailable surface AI on each ITN type after 20 washes may better explain the discrepancy in the chemical analysis and bioefficacy results with both pyrethroid-CFP ITNs after washing. Longitudinal field studies evaluating the durability of the insecticidal activity of PermaNet® Dual over several years under user conditions are advisable.

## Conclusion

In this laboratory study, we showed that PermaNet® Dual fulfilled WHO efficacy criteria against susceptible mosquitoes in laboratory bioassays and has potential to improve control of pyrethroid-resistant malaria vectors. These findings confirm those of recent experimental hut studies demonstrating that PermaNet® Dual represents an additional option within the highly effective pyrethroid-CFP net class for control of malaria transmitted by pyrethroid-resistant mosquitoes.

## List of abbreviations

AI: Active Ingredient
CFP: Chlorfenapyr
CRA-W: Centre Walloon de Recherches Agronomiques
cRCT: Cluster-randomised controlled trial
CREC: Centre de Recherche Entomologique de Cotonou
GLP: Good Laboratory Practice
ITN: Insecticide-treated net
LSHTM: London School of Hygiene & Tropical Medicine
OECD: Organisation for Economic Co-operation and Development
PBO: Piperonyl butoxide
PQT/VCP: Prequalification Unit Vector Control Product Assessment Team
RSD: Relative standard deviation
SOP: Standard Operating Procedure
WHO: World Health Organisation
WHOPES: World Health Organisation Pesticide Evaluation Scheme

## Availability of study materials

All aggregated datasets used and/or analysed during the study are provided as supplementary information. The full disaggregated datasets used and/or analysed during the current study are available from the corresponding author on reasonable request.

## Authors’ contributions

CN designed the study, acquired funding, supervised the project and prepared the final manuscript. TS supervised the laboratory bioassays, analysed the data, prepared the graphs and prepared the draft manuscript. BN performed the laboratory bioassays while DT performed the susceptibility bioassays. VA ensured compliance of the study to principles of Good Laboratory Practice. All authors read and approved the final version of the manuscript.

## Acknowledgments

We thank Melinda Hadi of Vestergaard Sàrl and Dr. Eleanore Sternberg (previously of Vestergaard Sàrl) for providing the PermaNet® nets and Olivier Pigeon and Patricia De Vos of CRA-W for performing the chemical analysis of the net pieces. We also thank the staff of CREC/LSHTM (Renaud Govoetchan, Josias Fagbohoun, Estelle Vigninou etc.) for their assistance and Ms. Imelda Glele and Danielle Apithy for administrative and logistics support.

